# HSP-4/BiP expression in secretory cells is regulated by a lineage-dependent differentiation program and not by the unfolded protein response

**DOI:** 10.1101/388272

**Authors:** Ji Zha, Jasmine Alexander-Floyd, Tali Gidalevitz

**Author notes:** Corresponding author. Address correspondence to T.G.

## Abstract

Differentiation of secretory cells leads to sharp increases in protein synthesis, challenging ER proteostasis. Anticipatory activation of the unfolded protein response (UPR) prepares cells for the onset of secretory function by expanding the ER size and folding capacity. How cells ensure that the repertoire of induced chaperones matches their post-differentiation folding needs is not well understood. We find that during differentiation of stem-like seam cells, a typical UPR target, the *C. elegans* BiP homologue HSP-4, is selectively induced in alae-secreting daughter cells, but is repressed in hypodermal daughter cells. Surprisingly, this lineage-dependent induction bypasses the requirement for UPR signaling, and instead is controlled by a specific developmental program. The repression of HSP-4 in hypodermal-fated cells requires a transcriptional regulator BLMP-1/BLIMP1, involved in differentiation of mammalian secretory cells. The HSP-4 induction is anticipatory, and is required for the integrity of secreted alae. Thus, differentiation programs can directly control a broad-specificity chaperone that is normally stress-dependent, to ensure the integrity of secreted proteins.

## Introduction

Cellular identity is largely defined by the proteins expressed in the cell, or cellular proteome. While the composition of cellular proteomes is determined by gene expression, their functionality depends on successful folding, localization, and functional maintenance of expressed proteins. During cellular differentiation, rapid onset of new protein synthesis challenges the proteostasis, and may result in the production of dysfunctional proteins and folding stress, if not matched by corresponding increases in required chaperones (Kikis, Gidalevitz, & Morimoto, 2010). This is especially important for professional secretory cells, as they may produce large quantities of a particular secreted protein (Itoh & Okamoto, 1980; Logothetopoulos & Jain, 1980), making them extremely sensitive to the folding stress in the endoplasmic reticulum (ER). To accommodate the anticipated increase in newly synthesized proteins, the capacity of ER proteostasis network is expanded during differentiation through activation of the ER stress response (Wu & Kaufman, 2006), known as the unfolded protein response, or UPR (Walter & Ron, 2011).

In addition to expanded capacity, the specificity of proteostasis netwforks may also need to be modified during differentiation, since different secreted proteins may require different chaperones for their biogenesis (Nagai et al., 2000; Wanderling et al., 2007). An early study of differentiation of a B-cell line into antibody-secreting cells showed that while expression of the majority of ER proteins, including the HSP70-family chaperone GRP78/BiP, increased in proportion to the expansion of ER size, a small subset of ER proteins was preferentially upregulated, resulting in their increased local concentration within the ER (Wiest et al., 1990). This uncovered two aspects of the ER proteostasis remodeling – (1) the general ER expansion, to match the increased biosynthetic demand and (2) selective upregulation of some chaperones, which in this case were likely required to support immunoglobulin folding and secretion (Wiest et al., 1990). How this selective upregulation is achieved, and whether it requires the UPR machinery, is not well understood.

The canonical UPR signaling includes IRE1/XBP-1, ATF-6 and PERK pathways, comprising the three branches of UPR. IRE1 and/or XBP-1 are essential for differentiation of many secretory cells such as plasma cells (Reimold et al., 2001; K. Zhang et al., 2005) and eosinophils (Bettigole et al., 2015), and for biogenesis of exocrine pancreas and salivary glands in mice (Iwawaki, Akai, & Kohno, 2010; Lee, Chu, Iwakoshi, & Glimcher, 2005). IRE1 is an ER transmembrane protein that, upon sensing folding stress in the ER, cleaves mRNA of a b-ZIP transcription factor XBP-1; the resulting active spliced form of XBP-1 controls expression of molecular chaperones and other ER biogenesis genes (Ron & Walter, 2007). Ectopic expression of spliced XBP-1 in cultured cells is sufficient to induce expansion of the ER size and cell’s secretory capacity, while deletion of *xbp1* gene in the mouse B-cell lineage prevents development of antibody-secreting plasma cells (Reimold et al., 2001; Shaffer et al., 2004). In fact, XBP-1, together with a transcriptional repressor BLIMP1, are the two regulators required for plasma cell differentiation (Shaffer et al., 2004; Turner, Mack, & Davis, 1994). The *xbp1* gene is repressed in resting B-cells (Reimold et al., 1996), and BLIMP1 relieves this repression upon B-cell stimulation, leading to upregulated *xbp1* transcription (Iwakoshi et al., 2003; Shaffer et al., 2004). Thus, plasma cell differentiation program directly regulates the UPR transcription factor responsible for the general increase in the secretory capacity. Indeed, activation of UPR during plasma cell differentiation appears to be in anticipation of increase in secretory load, rather than in response to proteostatic stress (Gass, Gifford, & Brewer, 2002; van Anken et al., 2003).

Compared with the general ER expansion, much less is known about the second aspect of the ER proteostasis remodeling during differentiation – upregulation of select chaperones to match the cell-type-specific folding needs. Many ER chaperones are expressed in cell- and tissue-specific patterns during development; however, there are only few examples where the basis of this cell selectivity is understood at molecular level. One known example is client-specific chaperones, such as a collagen chaperone HSP47, which is normally induced by heat stress, but during development is co-regulated with its client collagens by developmental transcription factors (Yasuda et al., 2002). Another known example is tissue-specific chaperones, such as cytosolic cardio-vascular small heat-shock protein HSPB7 regulated by the Myocyte enhancer factor-2 (MEF2) (Wales, Hashemi, Blais, & McDermott, 2014). Third is the induction of two stress-responsive ER chaperones, BiP and calreticulin, during cardiac development (Guo et al., 2001; Mao, Tai, Bai, Poizat, & Lee, 2006; Qiu et al., 2008). Unlike the client-specific HSP47 or the tissue-specific HSPB7, BiP and calreticulin are broad-specificity chaperones, required for the general housekeeping functions in the ER. The *Grp78* gene, which encodes BiP, is a canonical UPR target, whose promoter has been used to delineate the UPR signaling and to identify binding motifs for UPR transcription factors (Kokame, Kato, & Miyata, 2001; Yamamoto, Yoshida, Kokame, Kaufman, & Mori, 2004; Yoshida, Haze, Yanagi, Yura, & Mori, 1998). The induction of BiP during early embryonic heart development reflects cooperation between the UPR transcription factor ATF-6 and the cardiac-specific transcription factor GATA-4, which appears to bind *Grp78* promoter through the ER-stress element (ERSE) that is otherwise recognized by ATF-6 under stress conditions (Mao et al., 2006). Calreticulin expression during cardiac development was also dependent on the cardiac-specific transcription factors, although the involvement of UPR transcription factors was not examined (Guo et al., 2001; Qiu et al., 2008). Thus, outside of the few client- and tissue-specific chaperones, it remains unclear whether selective upregulation of ER chaperones during differentiation represents a subset of the generic UPR signaling, perhaps guided by cell-specific factors, or whether it can be directly controlled by the cellular differentiation program.

Here, we take advantage of the stereotypical timing and patterns of cell divisions and differentiation in *C. elegans,* to examine the regulation of a broad specificity chaperone, the BiP homologue HSP-4, during differentiation of dedicated secretory cells that secrete cuticular ridges called alae. *C. elegans* possesses two homologues of BiP, HSP-3 that is both constitutively expressed and stress-responsive, and HSP-4 that has very low basal expression in most cells but is strongly induced by UPR signaling (Kapulkin, Hiester, & Link, 2005; Shen et al., 2001).

Using the well-characterized transcriptional reporter phsp-4::GFP (Calfon et al., 2002), we find that *hsp-4* is selectively and transiently induced during differentiation of the stem-like seam cells into alae-secreting cells. Asymmetric divisions of seam cells produce anterior daughters that differentiate into hypodermal cells, and posterior daughters that continue stem-like divisions, but differentiate into the alae-secreting cells after the last division. *hsp-4* is induced only in these posterior daughters prior to their differentiation, in an anticipatory fashion. Unexpectedly, this *hsp-4* induction is neither dependent on the three canonical UPR signaling pathways – IRE1/XBP-1, ATF-6, and PERK, nor requires the known ER-stress elements in its promoter. Interestingly, we find that BLMP-1, a *C. elegans* homologue of the B-cell differentiation factor BLIMP1, represses *hsp-4* transcription in the hypodermal-fated anterior daughter cells. The non-UPR induction of HSP-4/BiP may be selectively required for the folding or secretion of a specific client(s) in alae-secreting cells, as indicated by abnormal alae structures and compromised barrier function of the cuticle when HSP-4/BiP induction is abolished. Our results demonstrate that a broad-specificity molecular chaperone that is a canonical UPR target can be selectively regulated by lineage-dependent differentiation signaling, independent of UPR pathways, to ensure the integrity of the secreted proteome and functionality of the cell post differentiation.

## Results

### *hsp-4* expression is activated in stem-like seam cells prior to their differentiation into alae-secreting cells

Although basal expression of the UPR-inducible BiP homologue HSP-4 is low in most tissues of *C. elegans,* the p*hsp-4::GFP* transcriptional reporter is visibly induced in unstressed animals in two tissues – spermathecae and the lateral seam. Both of these tissues are highly secretory. Because seam cells undergo stereotypical and well-characterized asymmetric stem-like divisions, and differentiate at defined developmental stages (Joshi, Riddle, Djabrayan, & Rothman, 2010), we used them to examine the regulation of BiP expression. During reproductive larval development, seam cells of V1-V4 and V6 lineages (Fig. 1A) undergo two types of divisions – one symmetric division early in the second larval stage (L2), that doubles the number of stem-like cells, and four rounds of asymmetric divisions (Joshi et al., 2010). The asymmetric divisions produce anterior daughters that differentiate and fuse with hypodermal syncytium after each cycle of divisions (Podbilewicz & White, 1994), and posterior daughters that retain their stem-like seam fate until the last asymmetric division (Fig. 1A). After the last division in L4, they exit the cell cycle, fuse with each other, and begin secreting proteins to make specialized cuticular structures, named alae (Johnstone, 2000; Sapio, Hilliard, Cermola, Favre, & Bazzicalupo, 2005). In addition to this normal developmental sequence, early L2 animals under certain environmental stress conditions can enter into an alternative developmental program known as dauer diapause, resulting in formation of non-feeding and long-lived dauer larvae (Golden & Riddle, 1984). During dauer development, the seam cells differentiate at the end of the pre-dauer L2 larval stage, known as L2d (Fig. 1A), and secrete the dauer-specific cuticle and alae (Sebastiano, Lassandro, & Bazzicalupo, 1991). This differentiation is not terminal, however, since triggering dauer recovery back to reproductive development results in these cells resuming their stem-like divisions (Karp & Ambros, 2012).

**Figure 1.**
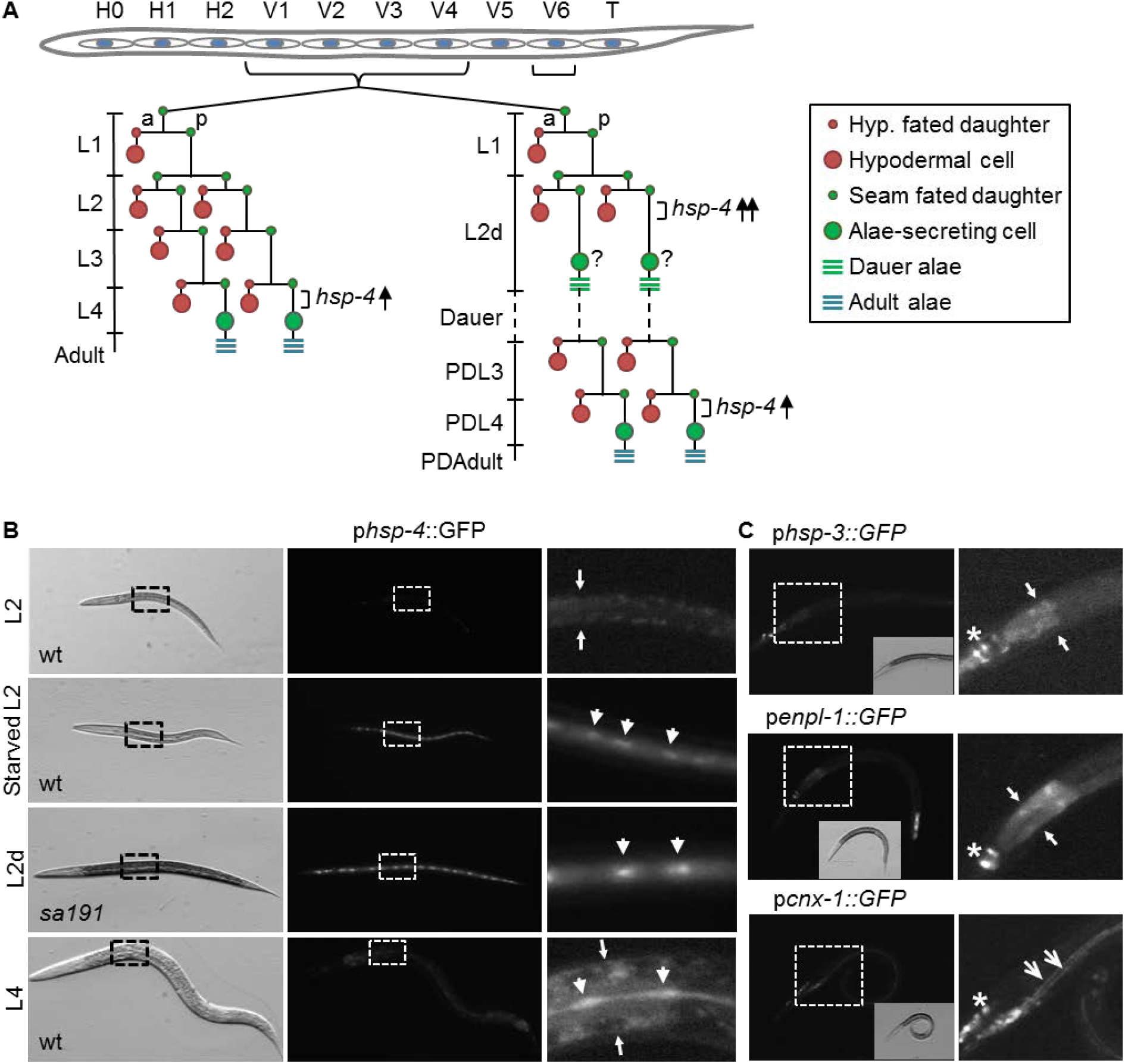
The UPR-inducible BiP homologue HSP-4 is selectively induced during differentiation of the stem-like seam cells. (A) Schematic diagram of cell divisions in V1-V4 and V6 seam lineages during *C.elegans* post-embryonic development. Positions and lineages of seam cells in L1 larvae are indicated in and above the worm outline. Left diagram corresponds to reproductive development, right – to dauer development. After most asymmetric divisions, anterior daughter cells (a) fuse with the hyp7 hypodermal syncytial cell, while posterior (p) daughters retain their stem-like seam fate. After the last asymmetric division in L4, posterior daughters initiate terminal differentiation into the alae-secreting cell, fuse with each other, and begin to secrete alae-constituent and other cuticular proteins (indicated by blue horizontal lines). Additionally, posterior daughters may undergo differentiation into alae-secreting cells during pre-dauer L2d stage, resulting in secretion of the dauer cuticle (green horizontal lines); these cells do not fuse and are not terminally differentiated (indicated by a question mark), as they resume asymmetric divisions in post-dauer (PD) animals. (B) Transmitted light and fluorescence micrographs of *C. elegans* expressing phsp-4::GFP at indicated developmental stages. L2d animals are *daf-28(sa191);* wt: wild type (N2). Images are taken at 120x magnification on stereo microscope, under same imaging conditions. Right panels show enlarged and (for L2 and L4) overexposed boxed areas. Small arrows points to intestine boundaries, arrowheads point to individual seam cells. (C) Micrographs, as in B, of *daf-28(sa191)* L2d animals carrying indicated transgenes. Right panels are enlarged and overexposed boxed areas, insets are transmitted light images. Small arrows point to intestine boundaries, open arrowheads to excretory cell, stars indicate fluorescence in the head.

We observed that p*hsp-4::GFP* reporter was visibly induced in seam cells during two developmental stages – weakly in the late fourth larval stage (L4), and strongly in L2 animals on starved crowded plates (Fig. 1B), while it was undetectable in other larval stages. Since starved L2 animals on crowded plates often initiate dauer program, and L4 and pre-dauer L2d are the only two stages in which seam cells exit cell cycle and differentiate (Fig. 1A), this expression pattern suggested a link between *hsp-4* induction and seam cell differentiation. To test this, we used a mutant allele (*sa191*) of an insulin/IGF-like protein DAF-28 that causes animals to enter the L2d pre-dauer stage even in the presence of food, and to remain in that stage for several hours (Klabonski, Zha, Senthilkumar, & Gidalevitz, 2016; Malone & Thomas, 1994). We observed a strong and persistent induction of the p*hsp-4::GFP* reporter in the seam cells of *daf-28(sal91)* animals at L2d stage (Fig. 1B).

Since *hsp-4* is a known target of UPR, we first asked whether this induction reflected activation of a generic UPR signaling, as in differentiation of B cells and other secretory cells. We tested available transcriptional reporters of three UPR target genes – hsp-3/BiP, *enpl-* 1/GRP94, and cnx-1/calnexin. While all three are known to be induced by ER stress in *C. elegans,* in IRE-1/XBP-1-dependent manner (Shen, Ellis, Sakaki, & Kaufman, 2005), none showed detectable induction in the seam cells of either L2d (Fig. 1C) or late L4 animals. Thus, the BiP homologue HSP-4 is selectively induced during differentiation of the stem-like seam cells.

We next thought to determine whether *hsp-4* gene expression was induced in anterior daughters, fated to differentiate into hypodermal cells, or in posterior daughters, fated to differentiate into alae-secreting cells after the last asymmetric division. To distinguish posterior daughters, we used a pegl-18::H1-mCherry transcriptional reporter for a GATA transcription factor EGL-18, which is necessary for adoption of the stem-like seam fate (Gorrepati, Thompson, & Eisenmann, 2013). This reporter is preferentially expressed in the posterior daughter cell after each asymmetric division, and equally in both stem-like daughters after the single symmetric division. We confirmed this known expression pattern in L1 and L2 animals (Fig. S1). However, *egl-18* reporter was not informative in pre-dauer L2d animals, as it was equally strongly expressed in both daughters, while *hsp-4* reporter was still only expressed in every other daughter cell (Fig. 2A). Since the asymmetric *egl-18* expression in larval stages is under control of the Wnt signaling pathway (Gorrepati et al., 2013), the difference in asymmetry between *egl-18* and *hsp-4* in L2d stage suggests that Wnt signaling does not determine the asymmetric induction of *hsp-4* during seam-cell differentiation.

**Figure 2.**
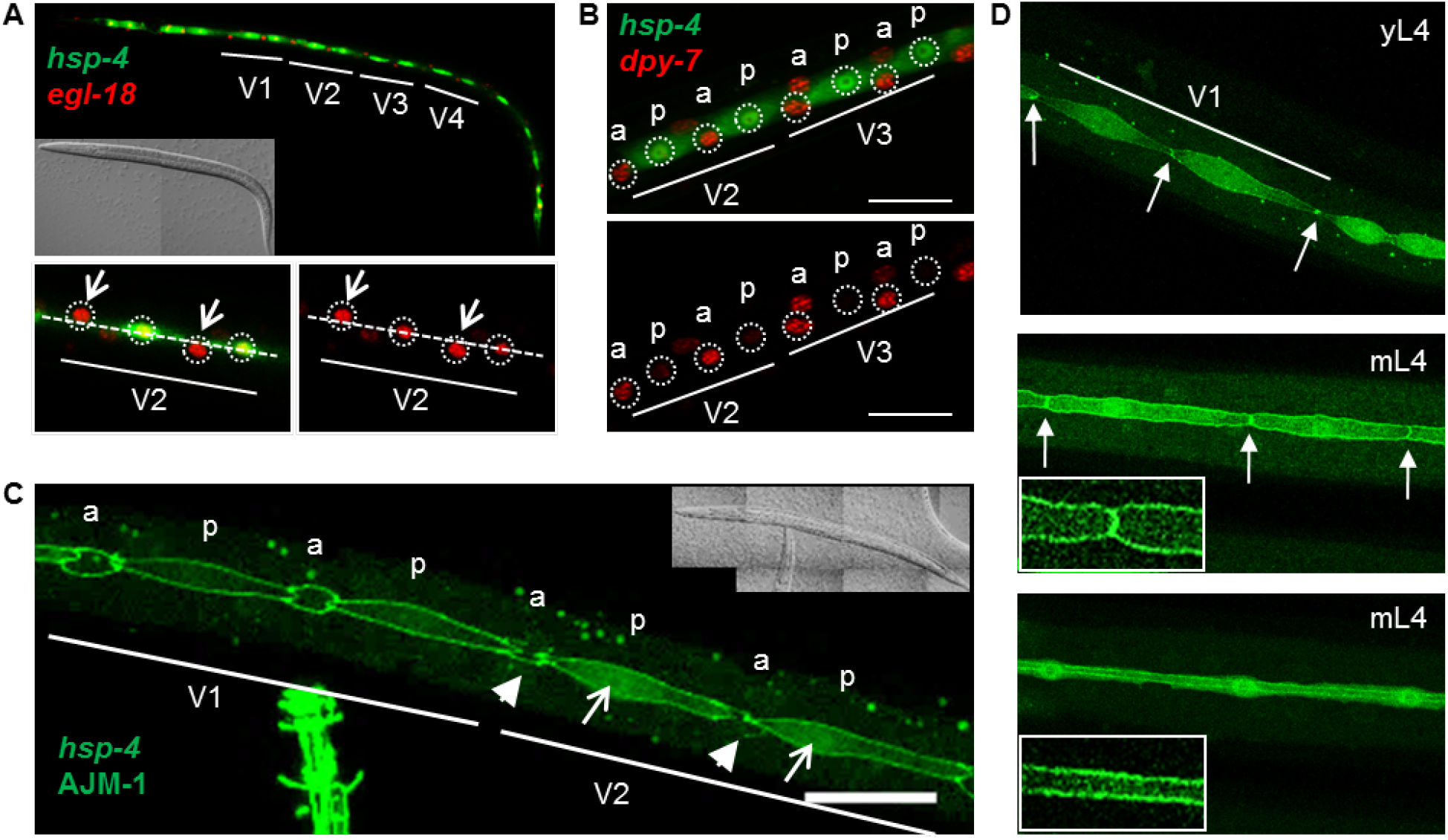
Expression of *hsp-4* is induced early during differentiation and only in the posterior daughter cells. (A) Confocal image of *daf-28(sa191)* L2d animal expressing phsp-4::GFP and *pegl-18::H1-* mCherry transgenes. Upper panel: projection of z-stack taken through the lateral hypodermis, lower panels: single plane image. Lower panels show close view of V2 lineage cells, either in both channels or in red channel only, with nuclei circled. Open arrows point to cells that appear migrating away from the seam center line (dashed line). (B) Close view of V2 and V3 lineage cells of a starved L2 wild type animal expressing p*hsp-4::GFP* and pdpy-7::HIS-24: :mCherry transgenes. The anterior (a) and posterior (p) daughters of V2 and V3 seam cell lineages are indicated, based on *dpy-7* expression. Z-stack projection, scale bar: 20μm. (C) Close view of V1 and V2 lineage cells of a *daf-28(sa191)* early L2d animal (39 hours post gastrula stage), expressing of phsp-4: :GFP reporter and AJM::GFP fusion protein. Open arrows point to already visible induction of *hsp-4* reporter in posterior daughters (p), which are outlined by the AJM::GFP protein; arrowheads point to the remnants of apical junctions of the differentiating anterior daughters (a). Scale bar: 20μm. (D) Seam cells of wild type animals expressing phsp-4::GFP reporter and AJM::GFP fusion protein. yL4, young L4 animal; mL4, mature L4 animal prior to (middle panel) or after (lower panel) seam cell fusion. Arrows point to the junctions between cells, which disappear after the fusion. Main panels: z-stack projections; insets: single plane images showing the AJM::GFP protein outlining either the cells prior to fusion (middle panel) or the seam syncytium (lower panel).

We noticed that hsp-4-negative/egl-18-positive nuclei in L2d seam cells appeared to be off the seam center-line (Fig. 2A), suggesting that these could be anterior daughters migrating towards their fusion with hypodermal syncytium. To test this, we used a transcriptional reporter of the cuticular collagen DPY-7, known to be expressed specifically in hypodermal cells (Gilleard, Barry, & Johnstone, 1997). *Pdpy-7*::HIS-24-mCherry reporter (Murray et al., 2012) was strongly expressed in *hsp-4*-negative cells and only weakly in *hsp-4*-positive cells (Fig. 2B). Thus, *hsp-4* expression is induced in the posterior seam cell daughters, which are fated to differentiate into alae-secreting cells.

### *hsp-4* expression is induced in stem-like seam cells in anticipation of their differentiation into alae-secreting cells

While the asymmetric expression pattern showed selective induction of the chaperone BiP/HSP-4 during differentiation, it did not allow for a distinction between anticipatory induction and that triggered by the post-differentiation increase in secretory load. To determine when during the last asymmetric division and differentiation *hsp-4* is induced, we used AJM-1::GFP protein that localizes to apical junctions in epithelial cells and outlines seam cell boundaries (Koppen et al., 2001). Immediately after the asymmetric division, AJM-1::GFP is present in both daughter cells. The differentiation of the anterior daughter and its fusion with the hypodermal syncytium results in loss of junctions between the differentiating cell and its neighbors, which can be detected by the loss of AJM-1::GFP signal (Harandi & Ambros, 2015). In contrast, posterior stem-like daughters continue expressing AJM-1::GFP until they differentiate, when they fuse and begin secreting proteins necessary for the formation of alae (Fielenbach et al., 2007). Based on AJM-1::GFP pattern in pre-dauer L2d animals, we determined that induction of *hsp-4* expression in posterior daughters happens already in the early stages after the last division, when anterior daughters just started to lose their boundaries and had not yet migrated away (Fig. 2C). This timing is consistent with anticipatory induction.

To further confirm the anticipatory nature of *hsp-4* induction, we examined its timing in *sa191* animals. Under normal growth conditions at 20°C, *sa191* animals that do activate the dauer program become L2d by 41 hours post gastrula. Because the dauer activation is only partial in these L2d animals, most of them (~70%) (Klabonski et al., 2016) return to reproductive development several hours later, instead of entering dauer (Malone & Thomas, 1994). Therefore, most of *sa191* animals do not complete the seam cell differentiation program and do not secrete dauer cuticle or form dauer alae. We found that the seam-cell – specific induction of *hsp-4* expression is readily detectable in 100% (n>100) of *sa191* animals that did enter L2d stage, assayed at 41 hours post gastrula, and is still present in the same animals at 46 hours post gastrula (see also control RNAi in Fig. 5A,B, n=63), after which time many animals return to reproductive development without secreting dauer cuticle or alae proteins.

To ask whether such anticipatory induction early in differentiation is peculiar to the pre-dauer stage, we examined seam cells in L4 larval stage. The last asymmetric division occurs around L3-to-L4 molt; alae-fated cells then differentiate and fuse at the end of L4 stage, prior to the onset of alae secretion (Joshi et al., 2010; Podbilewicz & White, 1994). The fusion is detectable by the change in AJM-1 ::GFP pattern from outlines of individual seam cells to the outline of the syncytium running along the body length of the nematode (Fig. 2D). Because L4 stage lasts nearly 10 hours at 20°C, we imaged the seam lineage in young L4s after the last asymmetric division, and in mature L4s, prior to and after the fusion. Expression of HSP-4::GFP reporter was evident already in the very young L4s, (Fig. 2D, upper panel), well before the fusion event, further supporting the anticipatory induction.

Finally, we asked whether downregulating expression of a major dauer alae protein in *sa191* mutant animals might decrease *hsp-4* induction in pre-dauer animals, which would argue against the anticipatory induction. *cut-1* codes for a cuticlin protein CUT-1, a disulfide-cross-linked protein that is expressed only in seam cells of pre-dauers and is a major component of the dauer alae (Sapio et al., 2005; Sebastiano et al., 1991). Although *cut-1* RNAi prevented dauer formation, as expected, it did not prevent *hsp-4* induction (Table 1). Collectively, these data show that *hsp-4* expression is selectively and anticipatory induced during differentiation of the alae-secreting cells.

**Table 1.** Genes targeted by RNAi. Genes tested in *daf-28(sa191);phsp-4::GFP* animals, using RNAi by feeding. Normal GFP pattern corresponds to induction in seam cells but low fluorescence in hypodermal cells. Blue shading indicates the presence of a binding peak in the *hsp-4* promoter in ModeEncode (pale blue=weak peak); green shading indicates known differential expression of NHRs in seam or hypodermal lineage (Gissendanner et al., 2004; Miyabayashi et al., 1999).

| Gene name | TF type or protein name | Stage for *hsp-4* binding | Normal GFP in seam cells? |
| --- | --- | --- | --- |
| *blmp-1* | homologue of BLIMP1 | emb, L1 | No |
| *nfyb-1* | subunit of NFY | L3, YA | Yes |
| F23B12.7 | CEBPZ (CCAAT/enhancer binding protein zeta) | YA | Yes |
| *elt-3* | GATA, controls hypodermal differentiation | emb, L1 | Yes |
| *mdl-1* | bHLH, similar to MAD, expressed in hypodermis, upregulated in dauer | L1 | Yes |
| *nhr-23* | NHR, regulates *hsp-4* in intestine | L3 | Yes |
| *nhr-77* | NHR | L4 | Yes |
| *nhr-66* | NHR |  | Yes |
| *nhr-72* | NHR |  | Yes |
| *nhr-74* | NHR |  | Yes |
| *nhr-75* | NHR |  | Yes |
| *nhr-80* | NHR |  | Yes |
| *nhr-81* | NHR |  | Yes |
| *nhr-82* | NHR |  | Yes |
| *nhr-89* | NHR |  | Yes |
| *cut-1* | dauer-specific cuticlin, required for alae formation |  | Yes |
| L4440 | RNAi empty vector |  | Yes |

### Developmental program signals *hsp-4* induction

Since *hsp-4* induction was strongest in pre-dauers and seemed to follow the initiation of the dauer signaling, we asked whether it was responding to a specific dauer-inducing signal. The decision to initiate dauer program *vs*. reproductive development is controlled by secretion from sensory neurons of two types of growth factors– insulins/IGFs and TGFβ. We found that *hsp-4* reporter was similarly induced in pre-dauer animals whether the dauer signaling was induced through insulin/IGF pathway *(daf-2(1370)* animals) or TGFβ pathway *(daf-7(e1372)* animals) (Fig. 3). Signaling through the transcription factor DAF-16/FOXO3, downstream of IGF receptor homologue DAF-2, was recently shown in *C. elegans* to have an impact on UPR (Henis-Korenblit et al., 2010). Thus, we also asked whether *hsp-4* induction was dependent on DAF-16. Animals bearing a hypomorphic allele *daf-16(mu86)* are dauer deficient; however those animals that did initiate pre-dauer program upon starvation/crowding had p*hsp-4::GFP* induction pattern indistinguishable from the wild type (Fig. 3). Finally, dauer induction requires the heat-shock transcription factor HSF-1 (Morley & Morimoto, 2004). *hsp-4* gene is heat-inducible, and *hsp-4* promoter has predicted HSF-1 binding sites (Fig. S2A). However, heat-shock response – deficient *hsf-1(sy441)* animals were still able to induce *hsp-4* in seam cells of starved L2s (Fig. 3).

**Figure 3.**
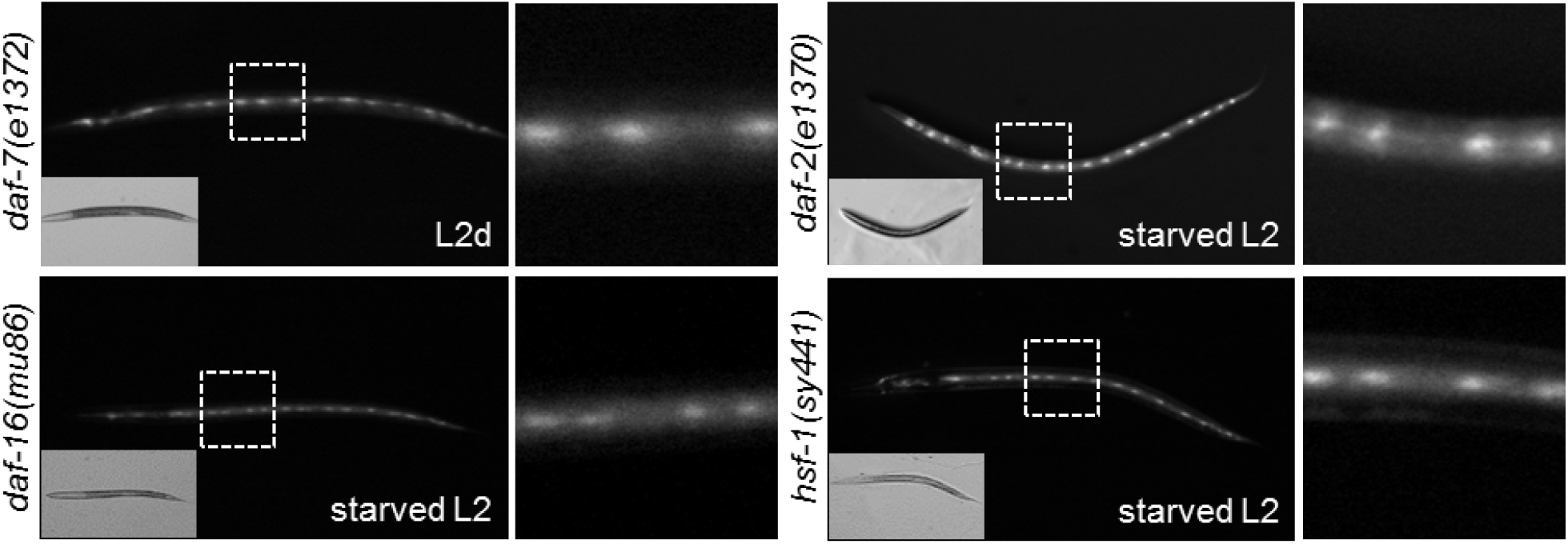
*hsp-4* induction during seam cell differentiation in pre-dauers does not depend on a specific dauer signaling pathway. Fluorescence micrographs of pre-dauer animals of indicated mutant strains, expressing p*hsp-4::GFP. daf-7(e1372)* mutant animals enter L2d stage at 20°C similar to the *daf-28(sa191)* animals; in other strains the pre-dauer stage was induced by starvation/crowding. Imaging as in Fig. 1B; right panels are close views of the boxed areas.

Since *hsp-4* induction appeared regulated by the dauer developmental program, we tested the most proximal signaling pathway in dauer development – steroid hormone signaling. As *C. elegans* possesses a large family of nuclear hormone receptors (NHRs), we chose among family members present in the RNAi library those that were previously shown to be differentially expressed in hypodermal or seam cells and/or affect alae production (Gissendanner, Crossgrove, Kraus, Maina, & Sluder, 2004; Miyabayashi, Palfreyman, Sluder, Slack, & Sengupta, 1999), regulate *hsp-4* in any tissue (MacNeil et al., 2015), or exhibited binding to *hsp-4* promoter based on modENCODE database (Van Nostrand & Kim, 2013) (Table 1). None of the NHRs tested by RNAi were required for the p*hsp-4::GFP* induction in seam cells. Besides NHRs, we tested two additional genes exhibiting *hsp-4* promoter-binding peaks in modENCODE database – *elt-3,* a GATA transcription factor regulating hypodermal differentiation, and *mdl-1,* a MAD-like transcription factor that is strongly upregulated in dauer stage (Wormbase), and found that neither is required for the induction of *hsp-4* reporter.

Together, our data show that HSP-4/BiP, but not other UPR-inducible chaperones, is selectively induced in stem-like seam cells prior to their differentiation into alae-secreting cells. The chaperone induction is anticipatory, linked to the last asymmetric division preceding differentiation, and is independent from Wnt signaling – the main asymmetry-determining pathway of the seam lineage. The HSP-4 induction is triggered by specific developmental programs – dauer entry and L4-adult transition, and is not dependent on any one individual dauer signaling pathway – insulin/IGF, TGFβ, or heat shock response pathways.

### *hsp-4* induction during seam-cell differentiation is independent from UPR signaling and does not require known ER stress elements in its promoter

Since induction of ER chaperones during differentiation of mammalian secretory cells depends on activation of some of the UPR pathways, we asked whether UPR pathways were required for the selective *hsp-4* induction during seam cell differentiation. We examined expression of p*hsp-4::GFP* reporter in seam cells of starved L2 animals deficient for each of the three canonical UPR pathways, by using loss of function alleles (Fig. 4A). These alleles were previously characterized as UPR-deficient and were shown to affect expression of *hsp-4* and other UPR target genes under both ER stress and basal conditions (Shen et al., 2005; Urano et al., 2002). Surprisingly, p*hsp-4::GFP* reporter was induced normally in seam cells despite inactivating mutations of *ire-1*/IRE1 or *xbp-*1/XBP-1, or deletions of pek-1/PERK or *atf-6*/ATF-6 (Fig. 4A). Mammalian ATF-6 and XBP-1 are both bZIP transcription factors, binding to similar DNA elements and capable of hetero-dimerization (Yamamoto et al., 2007). Genetic inactivation of each is well tolerated in *C. elegans*, but loss of both is larval lethal, due to degeneration of intestine (Shen et al., 2001). Thus, it is possible that they compensate for each other in the singly-deficient backgrounds. To test this, we used feeding RNAi to downregulate *atf-6* expression in *xbp-1*-deficient animals. To avoid the possible complications of combining feeding RNAi with starvation, we scored p*hsp-4::GFP* induction during differentiation of seam cells in L4 larval stage. All scored (n=20) xbp-1(zc12);*atf-6*(RNAi) animals had normal induction (Fig. 4B), despite being unhealthy with patchy coloration in their intestines, which indicated that RNAi treatment was effective (Shen et al., 2001).

**Figure 4.**
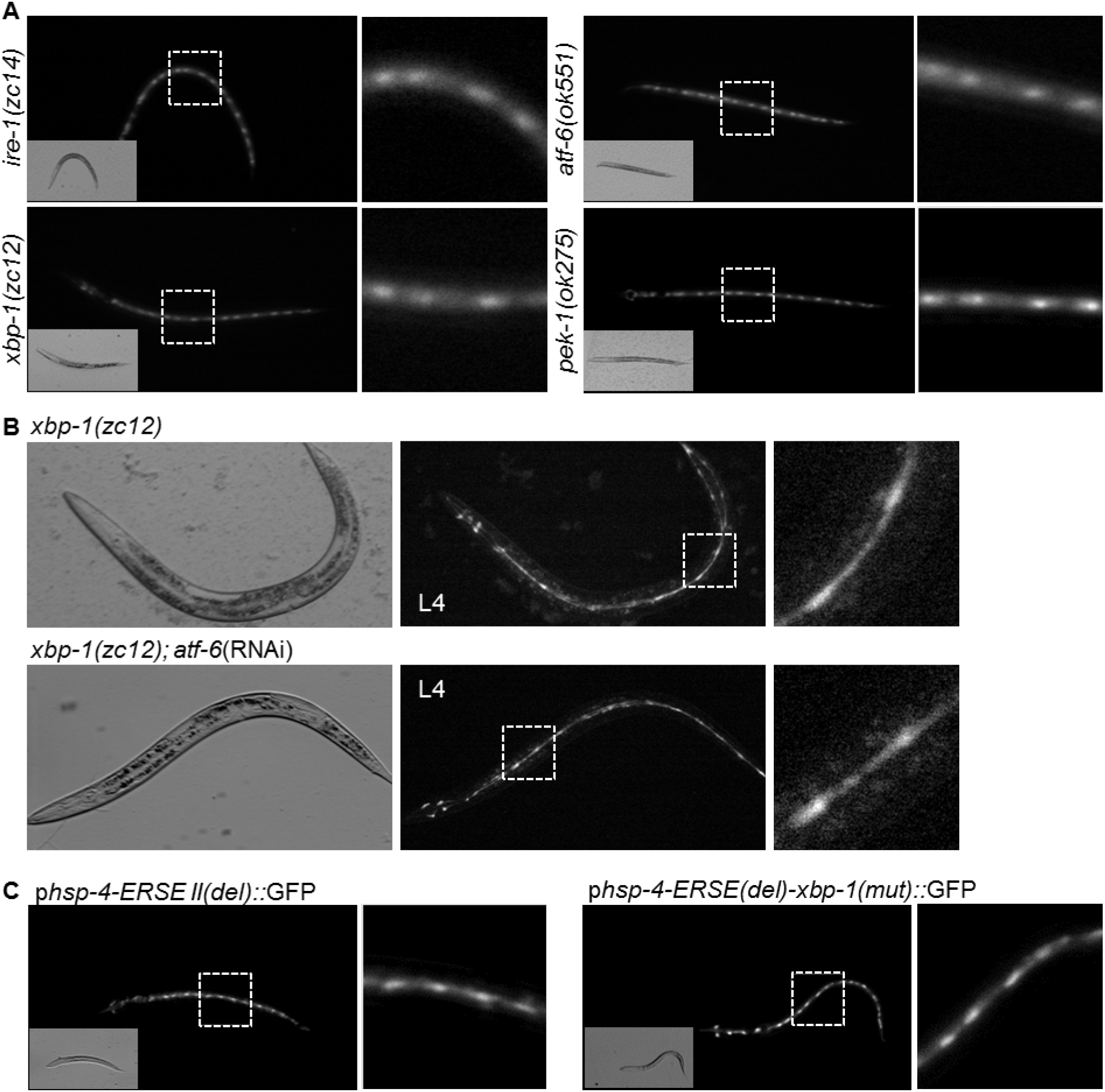
*hsp-4* induction during seam-cell differentiation is independent from UPR signaling. (A) Expression of *hsp-4* reporter in the seam cells of starvation/crowding induced pre-dauers of UPR-deficient mutant strains. All animals carry p*hsp-4: :GFP* transgene. (B) Combined loss of XBP-1 and ATF-6 transcription factors does not prevent *hsp-4* induction in differentiating seam cells. Upper panels, *xbp-1(zc12)* animals fed control RNAi (L4440 empty vector); lower panels, *xbp-1(zc12)* animal fed *atf-6* RNAi for two generations. (C) *hsp-4* reporter lacking either only ERSE II region (left panel) or both known ER stress elements (right panel) is still specifically induced in the differentiating alae-secreting cells.

We could not completely exclude the possibility that a small amount of ATF-6 protein was still expressed in RNAi-treated *xbp-1(zc12)* animals. To address this, we thought to mutate the ER stress elements in the promoter of *hsp-4* reporter. *hsp-4* promoter was previously found to contain two ERSE-II–like elements and a putative XBP-1/ATF-6 (CRE-like) element (Shen et al., 2001) (Fig. S2A). The *hsp-4* ERSE-II–like elements, ATTGG-N(6)-CCACA, show some deviation from ERSE-II consensus sequence ATTGG-N(1)-CCAC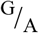, as well as from ERSE consensus CCAAT-N(9)-CCAC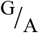, where CCAAT or ATTGG is a recognition site for the transcription factor NF-Y, while CCAC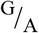 is recognized by XBP-1 or ATF-6 (Kokame et al., 2001; Yoshida et al., 1998). In the *hsp-4* promoter, the two ERSE–II–like elements and their flanking regions contain perfect reverse-complementary sequences, such that the region containing these elements, from residue (−584) to (−742), can form a highly stable stem-loop structure (Fig. S2A). Because of this unusual arrangement, we chose to delete, rather than mutate, this region. We found that deletion of the ERSE-II–like – containing region did not prevent the induction of *hsp-4* reporter in differentiating seam cells (Fig. 4C). To further confirm this, we downregulated two CCAAT/enhancer-binding proteins that bind *hsp-4* promoter, based on ModeEncode database: NFYB-1, a worm homologue of the beta subunit of NF-Y, and F23B12.7, a putative homologue of CEBPZ/CHOP, and found that neither RNAi prevented *hsp-4* induction in seam cells of starved L2s (Table 1).

The second ER stress element between nucleotides (−243) and (−269) is located on the reverse strand (Fig. S2A), and contains the TG<u>ACGT</u>GT XBP-1/ATF-6 (CRE-like) element, with the core XBP-1 motif underlined. We mutated this element to gGggGTGT (mutated residues in lower case) in the promoter with deleted ERSE-II–like region, thus eliminating both types of the known ER stress elements in this promoter (Shen et al., 2001). In agreement with the lack of effect from deleting UPR transcription factors, elimination of ER stress elements from *hsp-4* promoter did not prevent its induction in posterior daughter cells during seam cell differentiation (Fig. 4C). Surprisingly, this double-mutant promoter was still responsive to induction by ectopically overexpressed spliced XBP-1 (XBP-1s, in neurons, data not shown). It is possible that additional binding sites, distinct from the known XBP-1 site, exist in this promoter, or that XBP-1s activates the mutant promoter through interaction with another transcriptional regulator.

However, because of the data from *xbp-1*(zc12);*atf-6*(RNAi) animals (Fig. 4B), and the lack of induction of other UPR-target genes (Fig. 1C), we favor the conclusion that induction of *hsp-4* expression during differentiation of the seam cells is independent of the three canonical UPR branches.

### BLMP-1, the *C. elegans* orthologue of B-lymphocyte-induced maturation protein 1 BLIMP1, represses HSP-4/BiP induction in the anterior daughters after the terminal division

In addition to the UPR transcription factor XBP-1, the transcriptional regulator BLIMP1 is known to be involved in differentiation of many secretory cell-types in mammals, as well as in promoting and maintaining stem cell identity (Hohenauer & Moore, 2012). The *C. elegans* orthologue, BLMP-1, is necessary for formation of both adult and dauer alae (Horn et al., 2014). Interestingly, the seam cell divisions themselves are normal in *blmp-1* mutants, suggesting that it only contributes to the alae-secreting cell fate (Huang et al., 2014). Thus, we thought to determine whether BLMP-1 has a role in regulating *hsp-4* induction during differentiation of the stem-like posterior daughters into alae-secreting cells. Examination of modENCODE data showed a strong binding peak for BLMP-1 on *hsp-4* promoter (Fig. S2B). This is likely to represent a true binding peak, for two reasons: first, this site does not overlap with the extreme highly occupied target (xHOT) regions, which represent redundant and likely non-specific binding of multiple transcription factors (Araya et al., 2014). Second, we identified a sequence homologous to the known interferon regulatory factor (IRF) binding site, near the XBP-1/ATF-6 (CRE-like) element (Fig. S2A). This sequence, **TAA<u>GAAAG</u>C**TCTC**GAAAAGTC**, contains a perfect match to the IRF consensus sequence GAAA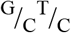 found in the MHC class I promoter (underlined), and partial matches (in bold) to interferon-stimulated response element, found in most interferon-inducible promoters (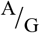NGAAANNGAAACT), as to the positive regulatory domains (PRD) element found in INF-β promoter (G(A)AA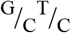GAAA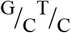) (Mamane et al., 1999). Because the mammalian BLMP1 is known to bind with high affinity to the subset of these elements containing GAAAG (Kuo & Calame, 2004), and the IRF consensus sequence in the *hsp-4* promoter contains this core sequence, we designate it as a putative BLMP-1-binding site (Fig. S2A).

To determine whether the developmental induction of *hsp-4* is dependent on BLMP-1 function, we downregulated *blmp-1* in *sa191 ;phsp-4::GFP* animals by RNAi. Under normal growth conditions, *sa191 ;phsp-4::GFP* animals begin entering L2d by ~30 hours post gastrula stage, and any animals that are not in L2d stage by 41 hours post gastrula will have bypassed the L2d entry and continued reproductive development. We found no effect of *blmp-1* RNAi on the p*hsp-4::GFP* reporter induction in all animals that had L2d morphology at 41 hours post gastrula (Fig. 5A). However, by 42 hours, *blmp-1* RNAi caused increased reporter fluorescence in seam cells of these animals, and by 46-47 hours, approximately half of *blmp-1* RNAi animals (n=81) exhibited induction of the reporter in the lateral hypodermis (Fig. 5A,B). None of the control RNAi animals (n=63) induced *hsp-4* reporter in the hypodermis. The induction level of p*hsp-4::GFP* reporter in the hypodermis of *blmp-1* RNAi animals was similar to that in seam cells, except for occasional one or few seam cells per animal that exhibited a much brighter further induction (Fig. 5).

**Figure 5.**
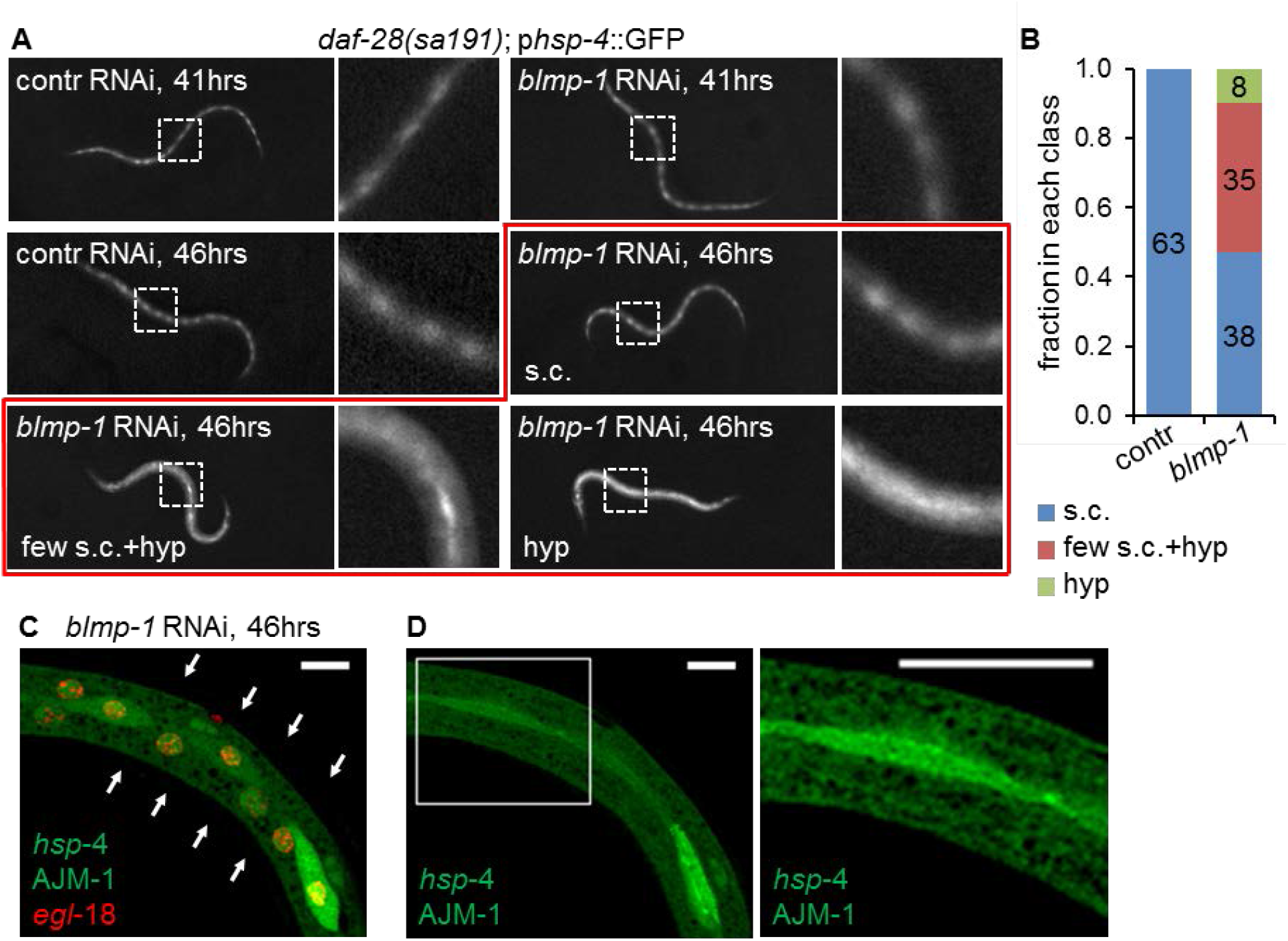
BLMP-1, the *C. elegans* orthologue of mammalian transcriptional regulator BLIMP1, represses *hsp-4* induction in the hypodermal lineage during differentiation of the alae-secreting cells. (A) Fluorescence micrographs of *daf-28(sa191)* animals at the early (41 hours post gastrula) and late (46 hours post gastrula) L2d stages. Downregulation of *blmp-1* expression by feeding RNAi results in strong induction of the *hsp-4* reporter in the lateral hypodermis (hyp) in a fraction of late L2d animals, as well as occasional hyper-induction in a few seam cells in addition to the induction in the hypodermis (few s.c.+ hyp). Animal with normal induction in seam cells are indicated as (s.c.). Imaging as in Fig. 1B. (B) Quantitation of the different *hsp-4* induction classes in animals shown in panel A. Animals that initiated the L2d stage were picked from indicated RNAi plates based on their morphology, using transmitted light only, and then scored for either normal *hsp-4* induction (s.c.), induction in the hypodermis (hyp), or induction in the hypodermis with hyper-induction in some seam cells (few s.c. + hyp). Numbers inside the bars indicate number of animals scored in each class, pooled from 3 independent RNAi experiments. (C, D) Confocal images of a representative animal with (few s.c. + hyp) pattern of *hsp-4* reporter induction following *blmp-1* RNAi. Images show phsp-4::GFP (green), AJM::GFP (green) and pegl-18::H1-wCherry (red). C: a single plane image, taken at a deeper plane through the lateral hypodermis, at the level of nuclei. *hsp-4* induction can be seen in the hypodermis; one hyperinduced seam cell is also visible. The arrows outline the animal’s body. D. Left panel shows a z-projection of the lateral hypodermis in the green channel only, same area as in C. Right panel is a close view of the boxed area, showing a single plane at the level of apical junctions. The AJM-1::GFP-outlined boundary between the seam cell and hypodermis is intact. Scale bar: 20μm.

We considered a possibility that the increase in fluorescence in hypodermal tissue resulted from re-distribution of the diffusible GFP protein from posterior seam cells to the hypodermis, if *blmp-1* RNAi caused defects in the seam-hypodermis boundary. However, examination of the AJM-1::GFP fluorescence showed that the boundary was intact in blmp-1(RNAi) animals with strong hypodermal *hsp-4* induction (Fig. 5C,D). Together, these data suggest that BLMP-1 normally represses *hsp-4* gene induction in the anterior daughters as they differentiate into hypodermal cells. Furthermore, the potentiation of *hsp-4* induction in some differentiating posterior daughters by *blmp-1* RNAi may indicate that BLMP-1 also regulates *hsp-4* gene in the posterior daughters at the early stages, thus explaining a hyper-response to a putative inductive signal in *blmp-1* RNAi animals.

We asked whether downregulation of *blmp-1* would result in induction of other ER chaperone genes. We examined same set of reporters as in Fig. 1C, and found that while *hsp-3,* encoding the second BiP homologue, was indeed weakly induced in seam cells of *sa191* L2d animals after *blmp-1* RNAi, the UPR targets enpl-1/GRP94 and cnx-1/calnexin were unaffected (Fig. S3). Thus, removal of BLMP-1-mediated suppression is not sufficient for the induction of general UPR target genes in the differentiating seam cells, and a BiP-specific inductive factor appears responsible for this developmentally-controlled expression of *hsp-4.*

### Loss of HSP-4/BiP expression interferes with structure and barrier function of the cuticle in adults and with alae formation in dauers

The logic of anticipatory and selective ER chaperone induction during differentiation would suggest that the upregulated chaperone is required for the specific secretory function of the resulting cell. Yet, BiP is considered to be a broad-specificity rather than client-selective chaperone, consistent with its global induction under folding stress conditions. We asked whether induction of *hsp-4/BiP* expression in differentiating alae-producing cells is important for the post-differentiation function of these cells, by examining the requirements for *hsp-4* for cuticular structure. A GFP fusion with a cuticular collagen COL-19 has been previously used to detect defects in the cuticle. This protein is expressed starting from late L4 stage, and is normally detected in evenly aligned circumferential pattern, as well as in the longitudinal linear structures of adult alae (Thein et al., 2003) (Figs. 6A and 7A). Downregulation of *hsp-4* by RNAi resulted in a disrupted circumferential pattern in young adults, such that 46% (n=13) of animals contained large gaps between the COL-19::GFP fibers overlaying the lateral hypodermis and those overlaying the ventral/dorsal hypodermis (Fig. 6B,C). In contrast, only 8% (n=12) of control RNAi animals had any gaps in cuticle. Furthermore, *hsp-4* RNAi caused occasional areas of disorganization of the longitudinal linear pattern, with COL-19::GFP being deposited in a “spaghetti”-like fashion in some animals (Fig. 6C). Similar gaps and disorganization are known to be caused by mutations in proteins involved in cuticle synthesis and molting (Cai, Phong, Fisher, & Wang, 2011; Meli, Osuna, Ruvkun, & Frand, 2010).

**Figure 6.**
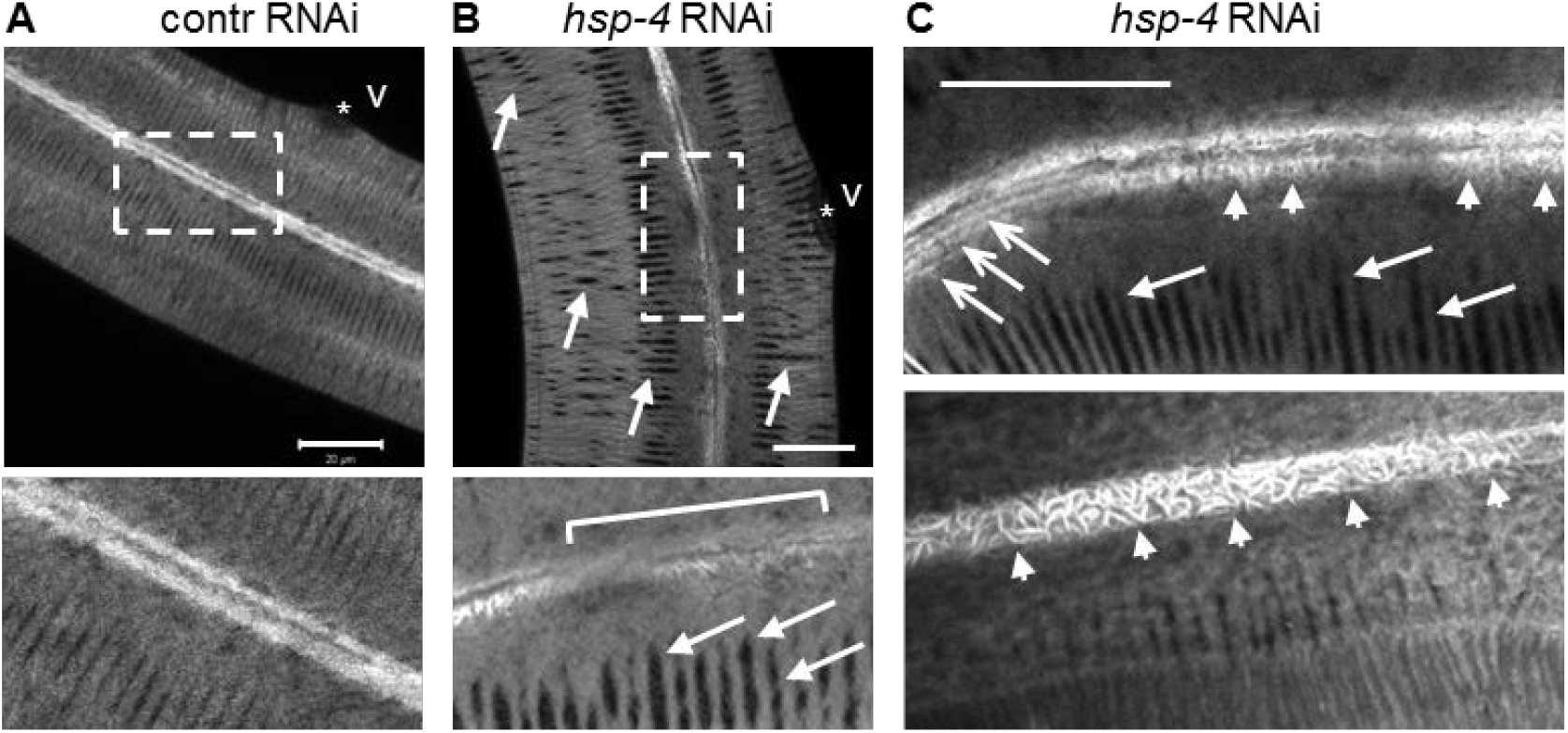
HSP-4/BiP is required for normal deposition of a cuticular protein. (A) Confocal image of COL-19::GFP protein in the hypodermis and alae of wild type (N2) young adult hermaphrodite, expressing COL-19::GFP protein, fed control RNAi (empty vector, L4440). COL-19: :GFP is deposited in the linear circumferential pattern in the hypodermis, and in linear longitudinal pattern underlying the alae structures. Star indicates position of the vulva (v). Lower panel: close view of the boxed region showing the linear alae pattern. Scale bar: 20μm. (B) Downregulation of *hsp-4* by feeding RNAi results in abnormal deposition of the COL-19::GFP protein, with large visible gaps in the circumferential pattern (arrows), and gaps and abnormal appearance of the longitudinal alae pattern (bracket). Scale bar: 20μm. (C) Examples of disrupted COL-19::GFP underlying the alae structures in adult hsp-4(RNAi) animals. Open arrows indicate area of normal longitudinal pattern, while arrowheads point to disorganized, “spaghetti-like” pattern of COL-19::GFP. Closed arrows point to the gaps in circumferential pattern. Scale bar: 20μm.

Because *hsp-4* and *hsp-3* genes share high degree of sequence homology, the *hsp-4* RNAi may target both genes. Thus, we confirmed the HSP-4 requirement for the cuticle using *hsp-4* deletion allele *gk514.* Unstressed *gk514* animals are phenotypically normal, and have normal dauer entry rates (Klabonski et al., 2016), presumably because of the stress-related role of HSP-4 and because the second BiP homologue, HSP-3, is functionally redundant with HSP-4 (Kapulkin et al., 2005). Yet, we found that deletion of *hsp-4* resulted in defects in COL-19::GFP deposition in young adults: the levels of COL-19::GFP over the lateral hypodermis overlaying the seam cells were strongly reduced, and the protein was absent in the longitudinal areas underlying the forming alae (Fig. 7A).

**Figure 7.**
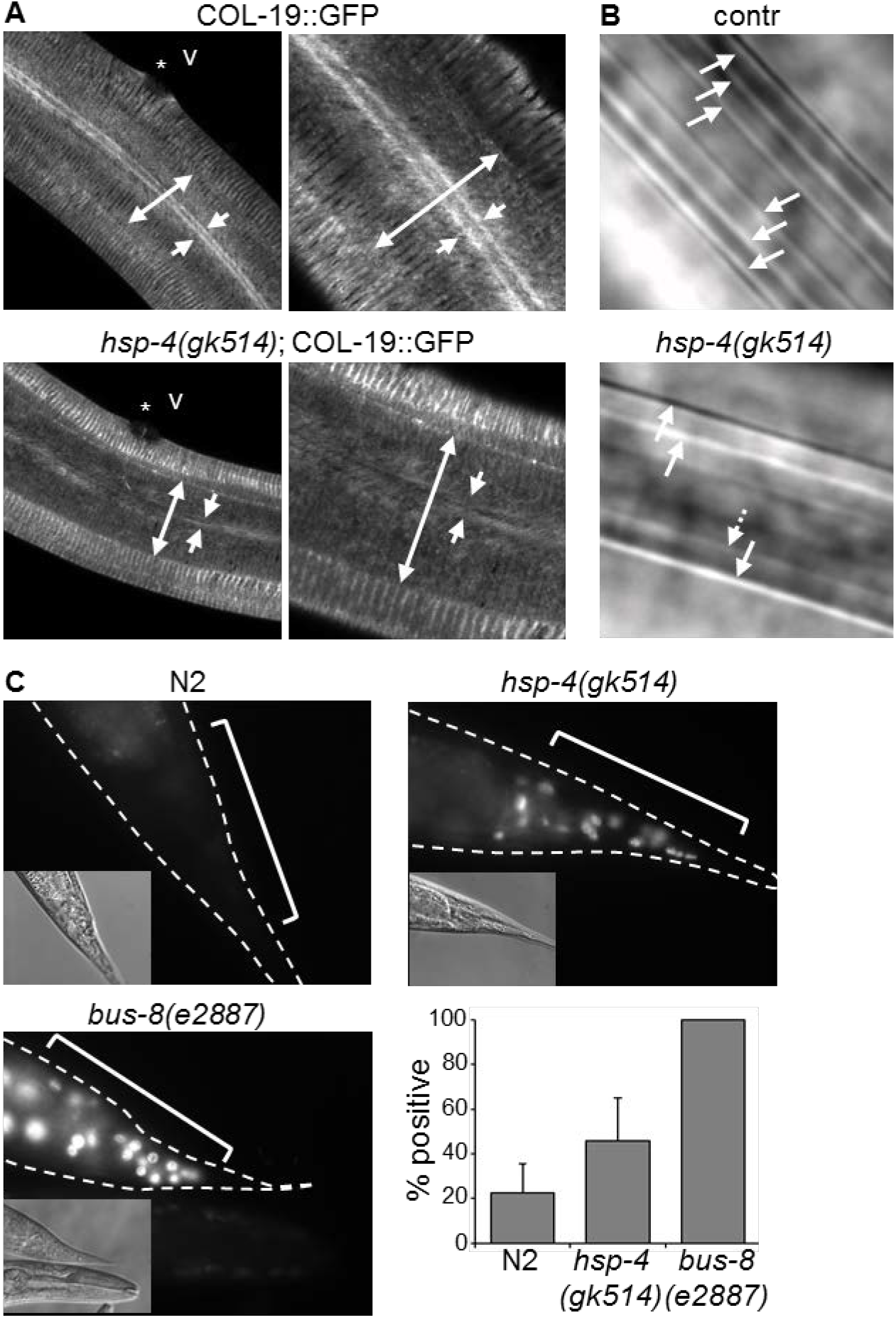
Deletion of HSP-4/BiP interferes with alae formation in dauers and barrier function of the cuticle in adults. (A) Genetic deletion of *hsp-4* results in decreased deposition of the COL-19::GFP protein in the lateral hypodermis. Double-headed arrows span the lateral hypodermis, overlaying the seam; arrows point to the location of the longitudinal COL-19 structures. (B) Dauer alae are abnormal in *daf-28(sa191)* dauers carrying *hsp-4(gk514)* deletion allele. Arrows indicate individual ridges, *daf-28(sa191);hsp-4(gk514)* dauer shown here has missing or flattened ridges. DIC images. (C) *hsp-4* deletion increases permeability of the cuticle to small molecules. Staining of nuclei in live animals treated with DNA-binding Hoechst dye reflects the degree of leakiness of the cuticle. Staining was performed as previously described (Hirani, Westenberg, Seed, Petalcorin, & Dolphin, 2016). Percent animals positive for the Hoechst staining is shown as bar graph, data are mean±SD of three independent experiments.

We next asked whether these structural defects affected the barrier function of the cuticle. Mutations that disrupt cuticle integrity cause it to become permeable to fluorescent dyes, such as Hoechst stain. We found that *hsp-4(gk514)* deletion caused increase in the cuticle permeability: although the degree of dye penetration was lower than that in the known leaky-cuticle *bus-8(e2887)* mutant strain (Partridge, Tearle, Gravato-Nobre, Schafer, & Hodgkin, 2008), twice as many *hsp-4(gk514)* than wild type animals took up the dye (Fig. 7C). Finally, because the *hsp-4* induction is strongest in pre-dauers, we examined the alae in dauer cuticles. As COL-19 is not expressed at these larval stages, we visualized the dauer alae by DIC microscopy. We found that deletion of *hsp-4* resulted in defective formation of dauer alae, with reduced number of ridges and visible gaps in the ridges in all examined dauers (n=5) (Fig. 7B). Thus, HSP-4 protein is required for the formation of the structurally and functionally intact cuticle, and this function of HSP-4 is not compensated for by HSP-3.

## Discussion

The importance of UPR signaling in the general expansion of ER biosynthetic capacity during differentiation of secretory cells has been firmly established. However, it is unclear whether UPR matches the repertoire of ER chaperones to the folding needs of specific secretory cell types. We find that during differentiation of the alae-secreting cells, induction of the stress-responsive *C. elegans* BiP homologue, HSP-4, bypasses the requirement for the canonical UPR signaling. Instead, HSP-4/BiP is induced by specific developmental programs – the dauer program or L4-to-adult transition. Interestingly, induction of HSP-4/BiP in the hypodermal-fated cells is repressed at the same developmental stage by a known transcriptional regulator of development, BLMP-1/BLIMP1, which also regulates differentiation of many secretory cell types in mammals. Importantly, induction of HSP-4 is not required for the differentiation of alae-secreting cells *per se,* but is essential for the secretory function of these cells post-differentiation.

Sharp increases in BiP expression are often interpreted to indicate activation of the UPR, and are thought to require ER stress elements in its promoter. However, under some conditions, the regulatory mechanisms differ from this expectation. For example, increased expression of BiP and other chaperones during acute-phase response in mice with bacterial infection was regulated by STAT3, which bound directly to *Gpr78* promoter (Ahyi et al., 2013). Even under conditions of ER stress, caused by limitation of specific folding resources, induction of the canonical UPR target proteins including BiP can be either dependent or independent of UPR signaling (Eletto et al., 2012). Thus, induction of *hsp-4* expression in our study may indicate action of a non-UPR transcription factor(s), specific to the differentiation of alae-secreting cells.

Another possibility is that a member of stress-responsive CREB/ATF family, other than ATF-6, is involved in regulating *hsp-4,* similar to the regulation of *Grp78* by ATF-4 during translation block (Luo, Baumeister, Yang, Abcouwer, & Lee, 2003), or involvement of OASIS in bone development (Murakami et al., 2009). However, mutating the CRE-like element in *hsp-4* promoter did not affect the pattern of its induction, making this possibility unlikely. Finally, it is possible that *hsp-4* is regulated during alae-secreting cell differentiation by a UPR transcription factor binding to an element other than the known ER-stress elements. This is an intriguing possibility, since we do see induction of the mutant *hsp-4* transcriptional reporter, lacking ER-stress elements, in neurons expressing spliced XBP-1.

The transcriptional induction of *hsp-4* gene during differentiation of alae-secreting cells appears to be under tight temporal control, and its timing supports anticipatory induction relative to its putative client protein(s). In L4 larvae, *hsp-4* induction precedes the fusion and differentiation by several hours, while in pre-dauers, we first detect the p*hsp-4::GFP* reporter fluorescence soon after the last asymmetric division, before the anterior daughters move away from the seam. Considering the time needed to accumulate the fluorescent signal to detectable levels, activation of *hsp-4* promoter is likely to occur even earlier. Even more strikingly, most *sa191* animals will have returned to reproductive development at 20°C instead of entering dauer, and thus will not have finished differentiation of the alae-secreting cells; yet, all *sa191* L2d animals strongly induce the *hsp-4* reporter. The anticipatory induction here parallels the regulatory logic of the UPR induction during differentiation of secretory cells in mammals (Gass et al., 2002; van Anken et al., 2003), even though it appears to bypass the UPR. It would be interesting in the future to understand whether this difference is an example of different organisms or even different cell-types using different routes to achieve the same goal – timely increase in the necessary chaperone, or whether it reflects the difference between the need for the generic expansion of ER capacity *vs* the need to match the chaperone repertoire to the cell-specific proteome.

HSP-4 induction is also not simply a consequence of asymmetric divisions, because it is not affected by the main seam asymmetry-determining pathway – Wnt signaling (Gleason & Eisenmann, 2010), and because *hsp-4* reporter is not induced after asymmetric divisions of these cells in other developmental stages. Together with the absence of a generic UPR in these cells, and with apparent independence of *hsp-4* induction from the canonical UPR signaling, these data suggest that the early differentiation program that determines the identity of the posterior daughter cell is able to directly regulate this chaperone. This phenomenon is similar to the recently reported regulation of some cytoplasmic chaperones by the myogenic transcription factor HLH-1/MyoD during embryonic muscle differentiation in *C. elegans* (Bar-Lavan et al., 2016). These chaperones, required for the myofilament formation, were found to have HLH-1-binding sites in their promoters. Similarly, the transcription factor Zf9 that regulates collagen-specific chaperone HSP47 in fibrotic tissues is capable of binding a collagen promoter (Yasuda et al., 2002). In these examples, the chaperones and their clients are regulated by the same transcription factor(s). While we do not know whether a similar regulatory logic applies to the developmental regulation of *hsp-4* gene, since transcription factors that specify the identity of the alae-secreting cells are unknown, our data do show that HSP-4 function is specifically required for the alae-secreting function of these cells post-differentiation.

Another aspect of the observed temporal control of *hsp-4* transcription is its repression in the lateral hypodermis by BLMP-1. Silencing of *blmp-1* resulted in *hsp-4* reporter expression in hypodermal cells, after the anterior, hypodermal-fated daughter cells fused with the hypodermal syncytium. Interestingly, this ectopic induction in blmp-1(RNAi) animals was observed only following the last division before alae-secreting cells are specified, but not during asymmetric seam cell divisions in other larval stages. The most facile explanation for such pattern of induction is existence of a positive inductive signal that is active in the entire seam lineage, and that temporally coincides with termination of stem-like divisions or onset of differentiation. In such a case, the combination of the inductive signal for *hsp-4* expression in the entire seam lineage of L2ds or L4s, with the repressive BLMP-1 function in the hypodermal-fated cells, should explain both the selective *hsp-4* induction in the posterior daughter cells of wild type animals and de-repression in the anterior daughters in blmp-1(RNAi) animals. Alternatively, the inductive signal may be specific to the posterior daughters as they assume the alae-secreting fate. In this case, *hsp-4* induction in the hypodermis upon *blmp-1* RNAi may reflect de-repression of a different factor that can induce *hsp-4* expression. Because our promoter sequence analysis suggests possible direct binding of BLMP-1 to *hsp-4* promoter, we favor the former scenario.

The dependence of the cuticle functionality on HSP-4 is surprising, since BiP is considered to be a broad specificity chaperone, capable of binding the majority of proteins that are folded in the ER (Blond-Elguindi et al., 1993; Flynn, Pohl, Flocco, & Rothman, 1991). In most *C. elegans* cells, HSP-4 protein is only expressed under folding stress conditions, further supporting the idea of non-selectivity of its function. Yet, its specific induction in the alae-secreting cell precursors, and the alae and cuticle defects seen with its deletion, suggest a unique cell-specific requirement for this chaperone. One possibility is that certain secreted proteins expressed in these cells require HSP-4, but not HSP-3, for their folding and secretion. Although HSP-4 and HSP-3 proteins are highly conserved and thought to be largely functionally redundant (Kapulkin et al., 2005), they are not identical, with 83% identity and 97% similarity in their peptide-binding domains. Another, less likely, possibility is that HSP-4 has a unique function in these cells, unrelated to its binding of unfolded proteins.

While the lack of *hsp-4* induction has clear negative consequences for the cuticle secretion, the functional importance of *hsp-4* repression by BLMP-1 in the hypodermal-fated cells is not immediately clear. Deletion of *blmp-1* was previously shown to cause defective formation of alae, and *blmp-1* -deficient animals have oxidative-stress sensitive cuticles and dumpy appearance (Hyun, Kim, Dumur, Schroeder, & You, 2016; L. Zhang, Zhou, Li, & Jin, 2012), indicating global cuticle defects. Because the *blmp-1* deletion is not cell-specific, we do not know whether these defects stem from functional deficiencies in the lateral hypodermis, where *hsp-4* is de-repressed in the absence of BLMP-1. However, it is possible that inappropriate induction of HSP-4 in hypodermal cells results in their decreased ability to secret proteins, because overexpression of a broad-specificity BiP chaperone under non-stress conditions and in the absence of high-affinity client may non-specifically stabilize folding intermediates and decrease rates of folding in the ER. Indeed, overexpression of BiP in CHO cells blocks secretion of a subset of proteins, while overexpression of its cytosolic counterpart, HSP70, causes developmental delays in *Drosophila* (Dorner, Wasley, & Kaufman, 1992; Feder, Rossi, Solomon, Solomon, & Lindquist, 1992). In addition to individual chaperones, ectopically increased UPR activity can be detrimental to animal development (Eletto, Eletto, Dersh, Gidalevitz, & Argon, 2014), and different tissues may have different tolerance levels (Taylor & Dillin, 2013).

The integration of developmental and stress signaling is emerging as an important contributor to multiple aspects of metazoan biology (Braakman & Hebert, 2013; Rutkowski & Hegde, 2010; Walter & Ron, 2011). UPR signaling pathways can be specifically activated in the absence of ER stress, for example by growth factor signaling or infections: IRE1 can be activated by internalized VEGF Receptor 2 through direct interaction (Zeng et al., 2013), while Toll-like receptors (TLR) in macrophages activate it by NADPH oxidase-dependent signal (Martinon, Chen, Lee, & Glimcher, 2010). Interestingly, the TLR-induced IRE1 activation does not result in chaperone expression or ER expansion, as would be expected from stress-activated IRE1, but rather promotes sustained production of inflammatory mediators (Martinon et al., 2010). Similarly, a canonical UPR transcription factor, ATF-6, and other members of CREB/ATF family respond to extracellular cues in osteoblasts and odontoblasts by regulating expression of collagens and other matrix-forming proteins (Kim et al., 2014; Murakami et al., 2009), presumably by interacting with cell-type-specific transcriptional machinery. Thus, physiological processes can not only induce the generic UPR activation, but can also trigger specific UPR pathways and, remarkably, control their outcomes. Our data show that, in addition, developmental signals can control the repertoire of induced chaperones directly, bypassing the UPR. Delineating the mechanisms integrating the physiological and stress signaling will thus be instrumental to further our understanding of the regulation of development, the pathogenesis of developmental disorders, and the mechanisms that maintain organismal homeostasis.

## Materials and Methods

*Strains and Genetics:*

Standard methods were used for worm culture and genetic crosses (Brenner, 1974). After crosses, strains were confirmed by PCR and restriction digest or sequencing. Animals were synchronized by picking gastrula-stage embryos from well-fed uncrowded plates.

The following strains were obtained from the *Caenorhabditis* Genetics Center (CGC):

SJ4005(*zcIs4*[p*hsp-4*::GFP]),
SJ30(ire-1(zc14) II; *zcIs4*[phsp-4::GFP]),
SJ17(*xbp-1(zc12*) III),
RB772(*atf-6(ok551*) X),
RB545(*pek-1(ok275*) X),
VC *1099(hsp-4(gk514)II),*
JT191(<u>daf-28(sa191)</u>V),
BC10514(*dpy-5(e907*) I; *sEx10514 [rCesT05E11.3::GFP + pCeh361]),*
BC10700(*dpy-5(e907)* I; *sEx10700 [rCesZK632.6::GFP + pCeh361*]),
RW11606(*unc-119(tm4063)* III; *stIs11606 [egl-18a::H1-wCherry + unc-119*(+)]),
SD1546(ccIs4251 I; stIs10166 [*day-7*p::HIS-24::*mCherry + unc-119*(+)].),
PS3729(*unc-119(ed4)* III; *syIs78*(AJM-1 ::GFP + *unc-119(+)*)),
CB1372(*daf-7(e1372*) III),
CF 1038(*daf-16(mu86*) I),
CB 1370(*daf-2(e1370*) III),
PS3551(*hsf-1(sy441)* I),
TP 12(*kaIs12* [COL-19:: GFP]),
CB6208(*bus-8*(*e2887*) X).
Strain expressing p*hsp-3::YFP* was a gift from the Morimoto lab (Northwestern University).
Wild type (N2) animals were a subclone of N2Bristol from the Morimoto Lab.

To generate pre-dauer L2d animals in *daf-28(sa191)* genetic background, appropriate strains were grown at 20°C under non-crowded/non-contaminated conditions on fresh plates seeded with OP50 *E. coli,* for at least 2 generations. 20-40 YA animals were then picked to fresh plates, and L2d larvae were picked among their progeny based on their morphology (Golden & Riddle, 1984; Klabonski et al., 2016): L2d animals are radially constricted, although to a lesser extent than dauers, they are larger than L2 but with germline morphology of L2, have a uniformly dark intestine, and exhibit slow pharyngeal pumping. Similar procedure was used for other developmental stages. To generate pre-dauer L2d animals by starvation/crowding, parents were placed on fresh plates seeded with OP50 *E.Coli* at 20°C, and plates were examined daily until there was no food left. Pre-dauers were imaged 1-2 days later.

*Transgenes Construction:*

p*hsp-4::*GFP-containing plasmid was obtained from Addgene (#21896). 54bp vector-derived region between the end of *hsp-4* promoter and start of GFP, which incidentally contained a PQM-1/DAE-like element, was removed using restriction enzyme PpuMI. To construct p*hsp-4-* ERSE II(del)::GFP transgene, 171bp of ERSE-II-like region was deleted using Q5 Site-Directed Mutagenesis Kit (NEB). To construct p*hsp-4-ERSE(del)-xbp-1(mut)::GFP,* the XBP-1/ATF-6 element in the *ERSE(del)* promoter was mutated from GATGACGTGT to GAgGggGTGT. All constructs were verified by sequencing (Macrogen USA).

The mutagenesis primers were:

ESERII_del_F: 5 ‘CGGGTCTCT AAGGAAAGGATTC3’
ESERII_del_R: 5’CCCAGTTGGACATCGGGTC3’
XBP_1_F: 5’CCTCTCCGATAAGTACACGTTGC3’
XBP_1_R: 5’GGGTGTATTAGTGCTGGAGAAATC3’

Transgenes were injected as a mix of 20 ng/μL plasmid DNA and 80 ng/μL sonicated salmon sperm DNA.

### RNAi

The RNAi clones were from Ahringer library. For RNAi experiments, animals were grown on 0.4 mM IPTG containing plates, spotted with designated RNAi bacteria, for one or two generations. For *blmp-1* RNAi, 15-20 L4 progeny of RNAi-treated *daf-28(sa191);phsp-4::GFP* parents were placed on fresh RNAi plates, gastrula-stage embryos picked, and L2d stage animals were scored 41-46 hours later. For *atf-6* RNAi, *xbp-1(zc12);phsp-4::GFP* animals were imaged at L4 stage. For *hsp-4* RNAi, COL-19::GFP deposition was examined in young adult animals. All experiments were repeated with a different population of animals 2-3 times. For all other RNAi experiments, pre-dauers were examined 1-2 days following exhaustion of bacteria. For *cut-1,* the plates were shifted to 25°C prior to exhaustion of bacteria, to enhance the dauer formation; animals were then examined daily, until dauers were seen on the control RNAi plates. No morphologically-normal dauers were found on *cut-1* RNAi plates.

*Microscopy:*

Confocal: animals were mounted on 2% agar pads, immobilized with sodium azide, and imaged with Zeiss LSM700 microscope, using 1.4NA 63x oil objective. Where indicated, 12 bit confocal z-stacks were reconstructed in ImageJ as 3D projections.

Stereo: animals were mounted as above, or immobilized by chilling on plates. Imaging was performed with Leica M205FA microscope and Hamamatsu Orca R2 camera, keeping magnification and intensity of fluorescence source (Chroma PhotoFluor 2) constant within each experiment.

## Acknowledgements

We thank Renee Brielmann (Morimoto lab) for transgene injections, and Drs. Yair Argon and Suraiya Haroon for comments on the manuscript. Some strains were provided by the CGC, which is funded by NIH Office of Research Infrastructure Programs (P40 OD010440). Confocal microscopy was performed at the Cell Imaging Center, Drexel University.

## Supplementary Figures

**Figure S1.**
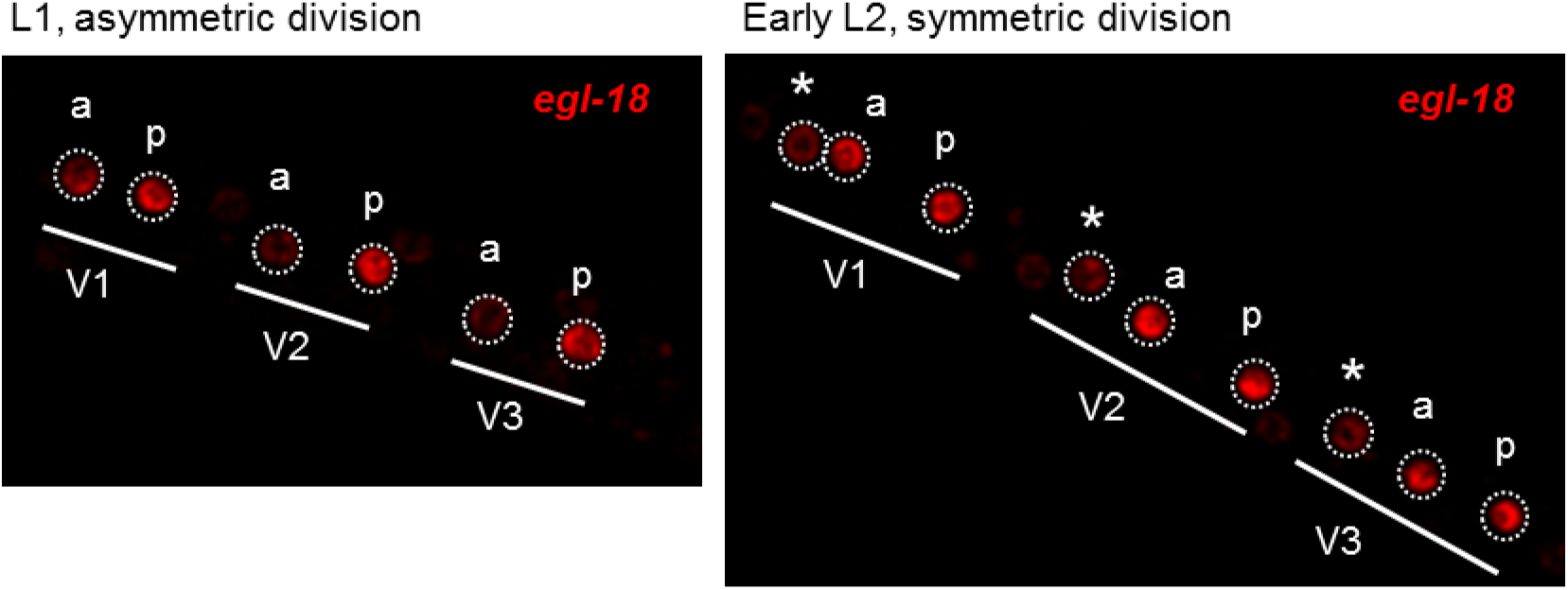
p*egl-18::mCherry* transgene identifies anterior and posterior daughters in L1-L2 *daf-28(sa191)* animals. Left panel, L1 animal (20 hrs post gastrula) shows differential expression of *egl-18* reporter in seam cells following the first asymmetric division. Anterior (a) and posterior (p) daughters are indicated. Right panel, early L2 animal (26 hrs post gastrula) following the symmetric division, with similar *egl-18* expression in anterior and posterior daughters; stars indicate anterior daughters from the previous round of division in L1.

**Figure S2.**
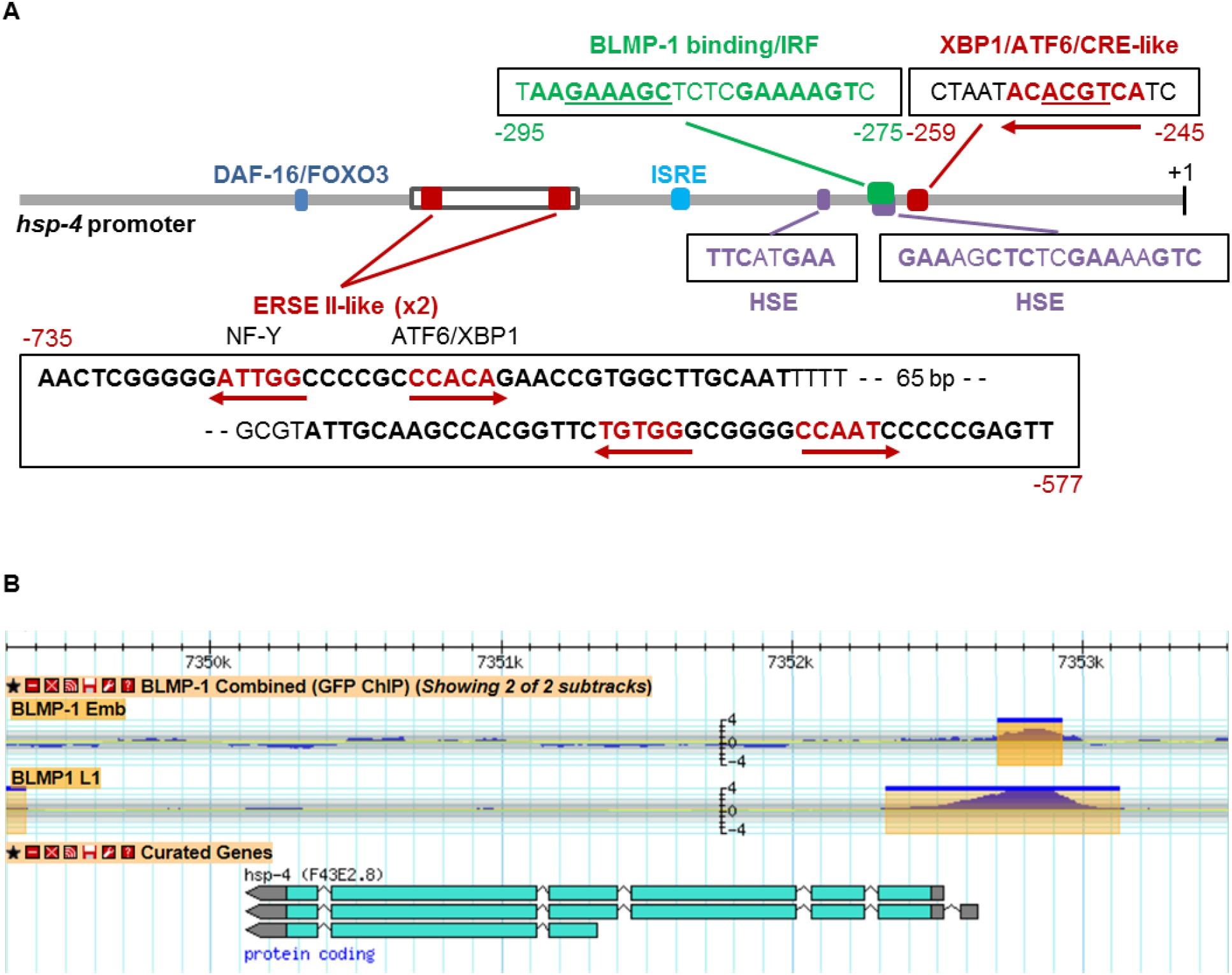
Regulatory elements in *hsp-4* promoter. (A) Schematic representation of the promoter used in p*hsp-4::GFP* reporter (grey line). Previously identified and putative regulatory elements/TF binding sites are indicated relative to the START site (+1). Corresponding sequences, their positions, and orientation relative to the sense strain are indicated. (B) Screenshot of the WormBase GBrowse image of BLMP-1 binding peak in *hsp-4* promoter, based on ModeEncode CHIP data.

**Figure S3.**
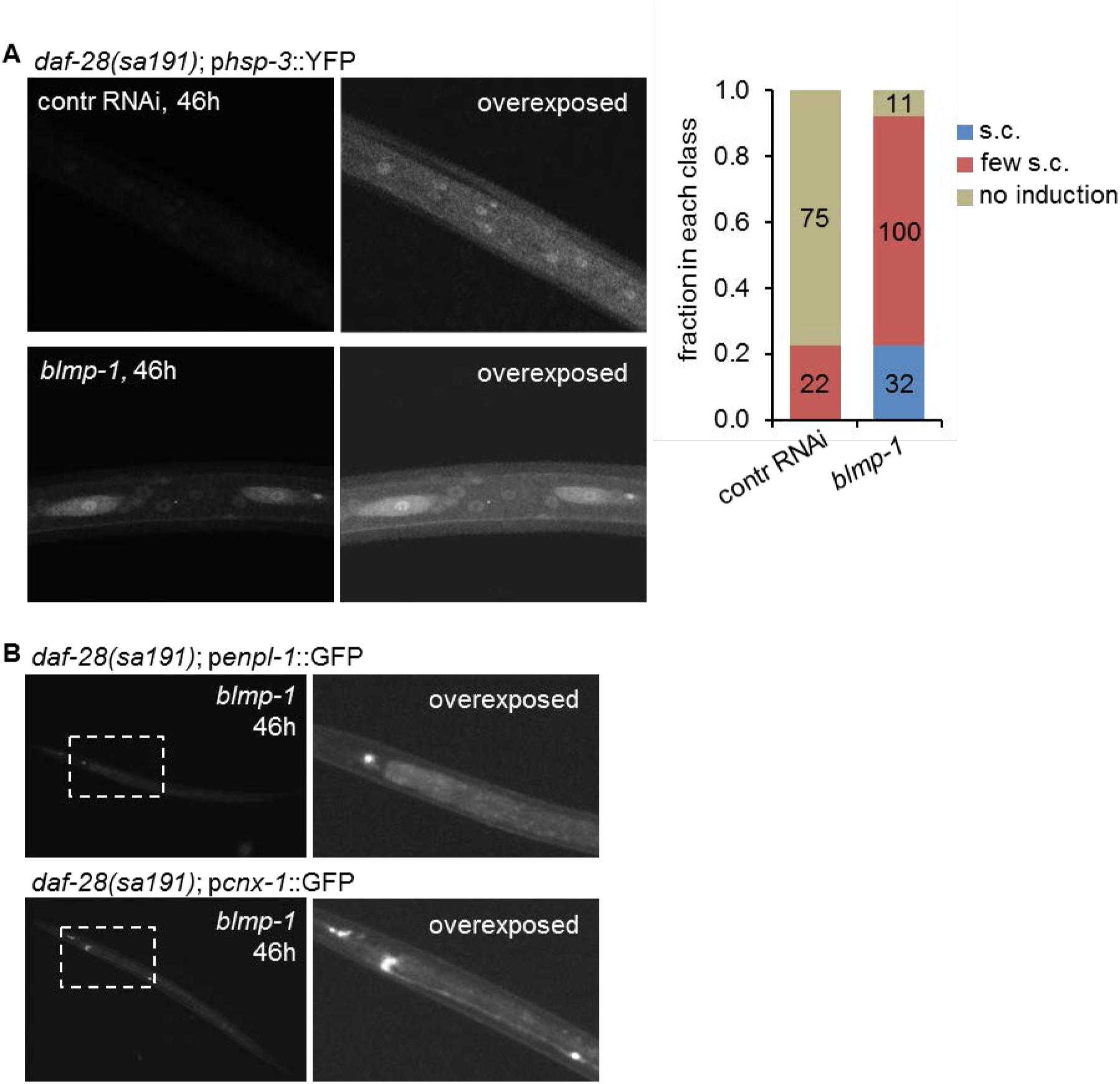
BLMP-1 represses both BiP isoforms, but not other UPR targets. (A) Downregulation of *blmp-1* results in mild induction of *hsp-3* expression in seam cells but not hypodermis of late L2d animals. RNAi and scoring as in Fig. 5, the expression classes scored were: induction in all seam cells (s.c.), induction in one or more but not in all seam cells (few s.c.), or no induction. (B) Downregulation of *blmp-1* did not result in induction in seam cells of two additional UPR-target genes, *enpl-1* and *cnx-1,* coding for *C. elegans* orthologues of GRP94 and calnexin, respectively.

